# New Insights from Old Data: Multimodal Classification of Schizophrenia using Automated Deep Learning Configurations

**DOI:** 10.1101/2020.11.02.364976

**Authors:** B Gagana

## Abstract

Schizophrenia is a heterogeneous cognitive disorder where clinical classification is challenging because of the lack of well-established, non-invasive diagnoses biomarkers. There is, hence, a need for objective systems that can classify Schizophrenia despite challenges such as overlapping symptomatic factors, diverse internal clinical manifestations, and complex diagnostic process leading to delayed treatment. Thus, experimentation with automated machine learning architectural frameworks (AutoML) is presented in order to handle multimodal Functional Network Connectivity(FNC) and Source Based Morphometry(SBM) features based on functional magnetic resonance imaging(fMRI) and structural magnetic resonance imaging(sMRI) components respectively. On evaluating the resultant AutoML models with respect to approximately 280 machine learning architectures on the Overall AUC metric, the former outperforms the latter despite remarkable limitations including complex high dimensional feature space with very little data.

## 1 Introduction

### 1.1 Schizophrenia

Schizophrenia is a mental fragmentation disorder[55] that is predominantly characterized by distorted perception of reality, hallucinations or delusions, and thought disorganization[16]. Diagnosis relies on the use of subjective processes of elimination due to the lack of well-established, non-invasive diagnoses clinical biomarkers[46], although there has been extensive interest in the medical community for its’ development[29][30] aimed at aiding diagnostic, prognostic and theranostic research[5].

The condition, however, with its elusive pathophysiology[39] is known to be surprisingly common^1^. The differential diagnosis of schizophrenia is however challenging specially while discriminating schizoaffective disorder from other closely related mental illnesses including clinical depression[46](specially in the premorbid and prodromal phases of preschizophrenia), bipolar I disorder[46][47] and attention-deficit/hyperactivity disorder(ADHD)[48]. Schizophrenics may thus receive multiple diagnosis based on appearance of symptoms which greatly influence treatment practices[31].

Schizophrenia is also associated with diverse internal clinical manifestations including abnormalities in anatomy of limbic-cortical systems[3], dysfunction in medial temporal (particularly hippocampal region) and lateral temporal lobe structures[40] and synaptic pruning[56]. However, at a neuronal level: the superior temporal gyrus appears to be less volumetric which negatively correlates with severity of corresponding abnormalities - hallucinations and thought disorders[2]. Schizophrenia is hence classified across a spectrum that ranges from attenuated schizoid and mild schizotypal personality to the most severe and debilitating forms[32].

Factors that potentially affect clinical assessments are:

a. Prolonged use of potent stimulants like Methamphetamines or hallucinogenic drugs like Lysergic acid diethylamide(LSD) that can impart schizophrenia-like symptoms[8].
b. Schizophrenia is known is to run in families[36] but there looms uncertainty concerning the degree of genetic contribution to the phenotypic variance of the disease[57]. These observations have led to scientific speculations that interactions between internal genetic components and external environments could trigger the disease[10].
c. Complications during pregnancy, childhood trauma, fetal growth retardation[58], inflammation of the brain, head injury, emotional consequences of imprisonment, and hypoxia[49] are also identified as activation points.
d. Research also further examines the role of external factors like employability and financial status of the patient as well as age and workload of the treating physician in the patient-physician relationship through the continuity of care (COC) metric[7].

To prevent any interference with diagnosis and subsequent treatment, diagnostic procedures rely on metrics including DUP (Duration of untreated psychosis) and DPIT (Delay in intensive treatment) which, in case of Schizophrenia could fail to account for the neurodevelopmental disturbances resulting in cognitive deficits before the onset of psychotic symptoms. The metrics measure a median of around twenty four weeks[35] which further highlight the need for early clinical or technological interventions[34].

### 1.2 Automated Machine Learning

With tremendous growth in the amount of data and complexity, designing machine learning systems is expertise intensive and tuning machine learning systems is time consuming. Automated machine learning (AutoML) attempts to automate these pipelines eliminating the confusion of whether certain models really perform better or were simply better tuned.

The generic workflow of a typical AutoML framework is as follows: The training data is fed into a system that automatically generates model architecture which has associated configurations that are tunable and hyperparameters which can be shipped for further predictions to be made on new data.

But the comparison of various AutoML frameworks and configurations is often tricky because[6]:

a. Blackbox systems in the background differ in their optimization methodologies, pipeline generation techniques, internal architectures and processing procedures while operating on similar data[13]. The results may also vary on other parameters including training time, memory and computational resources.
b. Reruns of the same configurations on data often yield different results. This tends to affect result reproducibility and complicate evaluation process.
c. Some frameworks automate the entire pipeline seamlessly while others tend to rely on external engineering techniques. In the latter case, performance evaluation is not a wholesome indicator of the framework’s capabilities. Across the machine learning pipeline, most of the AutoML tools have fairly similar processes with respect to data cleaning, data coding, metric selections but differ in their algorithm selection, parameter optimisation and post processing abilities.

Multiple surveys of automated machine learning frameworks focus on evaluating tools on various data[6][13][14][15]. But investigations are limited on the performance of various configurations of these tools on similar data. Although current surveys do help perform guided research to some extent, the need for trial and error evaluation of tools is not completely eliminated. There are no universal standardised configuration benchmarks and that necessitates trial and error evaluation in order to select the right tools and corresponding configurations on any given data. In this context, automated machine learning has not been explored extensively and prior research has shown that automated frameworks and tools do not necessarily perform better than machine learning algorithms[6]. This paper hence explores the use of automated machine learning (AutoML) on multimodal schizophrenia data to outperform classification algorithms.

This paper aims to study the architectures and corresponding performances of AutoML on multimodal data, with specific metrics particularly in healthcare. Also, as such, classification of individual subjects based on observed behavioral features[12] is more challenging than reporting group differences via voxel based approaches[1]. Some inconsistencies resulting from data representativeness heuristics[27] reflect the true reality of neurological research thus making this data a perfect challenge for AutoML.

## 2 Background and Related work

This section is intended to reflect on past literature in both biomedical research in Schizophrenia classification as well as in automated methodologies applicable to healthcare.

### 2.1 Current Clinical Detection Processes

Most of the defining attributes of schizophrenia are based on inferences drawn from self reported subjective experiences (along with but not limited to psychiatric status observation of family and personal history - evaluation of demeanor, appearance, thought process about delusions, hallucinations, and potential for violence or suicide) owing to the complexity of the disorder and the ambiguity in its diagnostic concept itself. The detection process involves a comprehensive medical examination with treatment regimen to reduce appearance of symptoms[42].

### 2.2 Exploration of AutoML for healthcare systems

The evolution of machine learning has contributed significantly to the healthcare industry through image detection, classification, and segmentation tasks but with respect to interest in mental health, there has been massive transformation specially with non-invasive neuroimaging[18]. For schizophrenia, in particular, machine learning methods[43] have been explored for diagnostics and classification from structural MRI[44] and functional MRI features[45], EEG signals[11] and resting EEG streams[4].

However, with respect to AutoML, the current research portfolio is sparse. One of the open benchmarks involving AutoML tools[6] suggests that:

i. AutoML does not perform better than untuned machine learning models(random forests).
ii. There is no AutoML system which consistently outperforms all others.

For blood transfusion in particular, H2O performs best as compared to other AutoML frameworks.

Another similar study[15] used AutoML to classify the following healthcare datasets:

a. Breast cancer(OpenML dataset ID 15) where ATM[26] seemed to perform the best(Average accuracy: 0.98474).
b. Diabetes(OpenML dataset ID 37) where hyperopt seemed to perform the best(Average accuracy: 0.7996).
c. Gene splice data(OpenML dataset ID 46) where hyperopt seemed to perform the best(Average accuracy: 0.9654).
d. Blood-transfusion data(OpenML dataset ID 1464) where RoBO seemed to perform the best(Average accuracy: 0.8076).
e. EEG eye state data(OpenML dataset ID 1471) where SMAC seemed to perform the best(Average accuracy: 0.9741).

## 3 Data Description

Owing to the adversities articulated in Section 1, there is a need to design automated processes that can classify schizophrenia but the technical challenge is to optimally combine the resting-state functional MRI and the gray-matter concentration information of structural MRI to automatically predict diagnosis labels and optimize performance. Schizophrenia is believed to be better characterized by the effective relationship between functional and structural data[27] which further assists in the understanding of pathology of the brain in general and specifically, contributes to the hunt for biomarkers.

### 3.1 Data Collection

The training data has 378 functional network connectivity features and 32 source based morphometry features with information on 144 subjects(75 healthy controls and 69 schizophrenic patients) acquired using a 3T Siemens Trio MRI scanner with a 12-channel head coil[27].

### 3.2 Data Preprocessing

SPM5 standard preprocessing technique ^2^ was employed to eliminate volumes leading to equilibration effects, image realignment, and correction of discontinuities in slice timings. Data was further spatially normalized, re-scaled to a mean of 100 and analyzed with spatio-temporal regression group independent component analysis (GICA)[27][60]. Independent component analysis further extracts artifacts embedded in the data from the preprocessed output using spatio-temporal regression[37]. The structural and functional data were then mapped into the space of a healthy control baseline dataset from which features were derived.

In the resultant data features, it was observed that there is little correlation between the structural and functional components but the groups amongst themselves are highly correlated.

### 3.3 Data Features

#### 3.3.1 Functional MRI: Functional Network Connectivity features

These functional feature modality values describe the overall connection levels between pairs of independent brain maps over time. The connectivity pattern between brain networks (also called synchronicity between brain areas) is mapped using the functional network connectivity features where the time courses indicate the activity level of the corresponding brain map at that point using the Pearson correlation values and hence, allows exploration of the neural interaction properties within the brain. Disruptive patterns are known to be indicative of schizophrenia[59]. The FNC matrix is symmetric and only the unique values are retained as features in this classification context[20].

**Figure 1:**
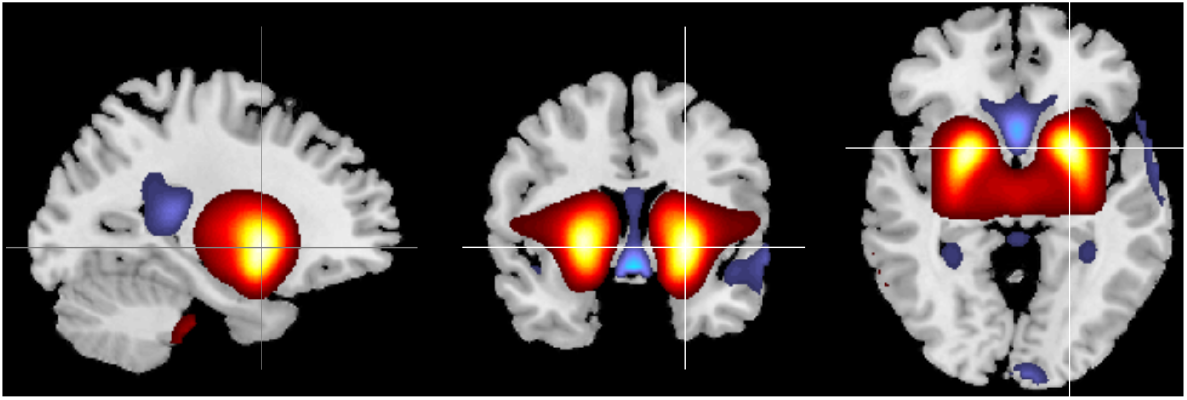
Visualization of FNC features

##### Observations

The data distribution of the FNC features is near-normal as shown in Figure 2.

**Figure 2:**
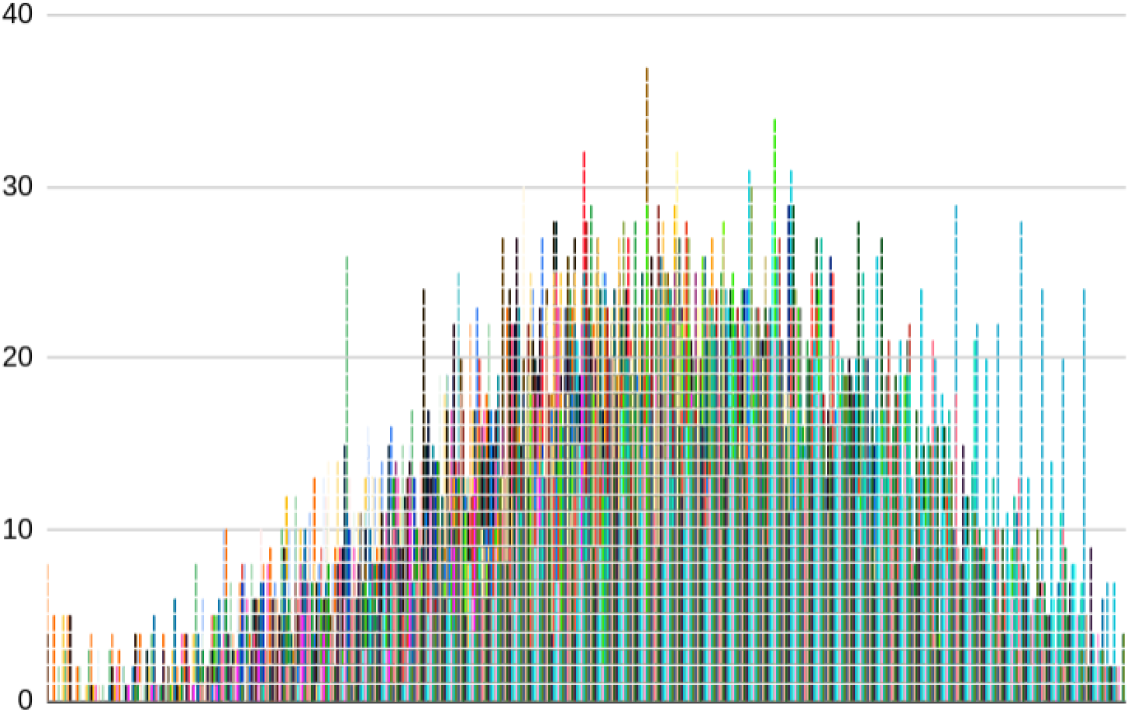
Visualisation of distribution of FNC features[20]

#### 3.3.2 Structural MRI: Source Based Morphometry features

Source based morphometry loadings[21] correspond to the standardised weights of all subjects with respect to the level of independent component analysis(ICA) brain maps derived from grey-matter concentration. The activity of the signal processing is further indicative of the computational power available in that region of the brain that is monitored along with the tissue density of grey matter, white matter and cerebrospinal fluid in the region[38][50][51]. A near zero structural modality feature value for given ICA derived brain maps is indicative that the brain regions in that subject are lowly present and hence, suggestive of schizophrenia[52][53][54].

##### Observations

The data distribution of the SBM features appears normal as shown in figure 4.

**Figure 3:**
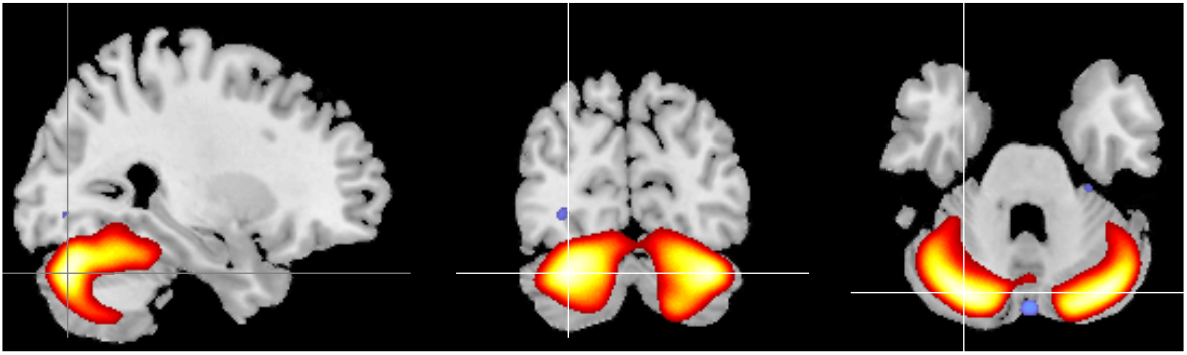
Visualization of SBM features[21]

**Figure 4:**
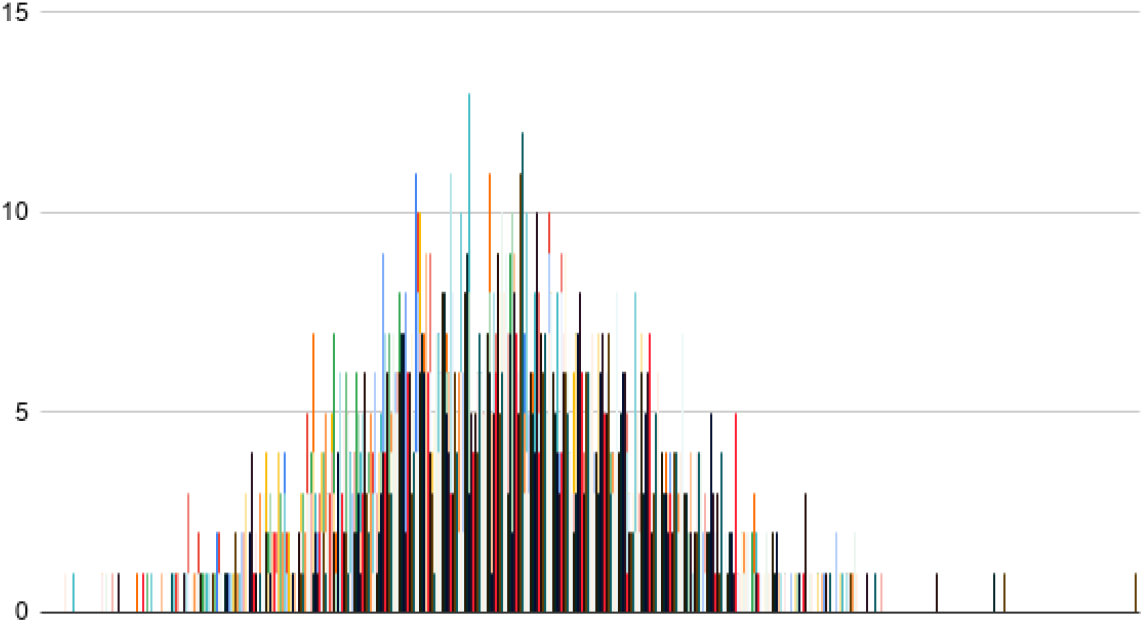
Visualisation of distribution of SBM features

## 4 Methodology

This section deals with two aspects:

a. Description of machine learning algorithms against which automated methods are compared and
b. The following open-source automated machine learning frameworks: Ludwig[22], H2O[23], AutoGluon-Tabular[24], MLBox[25], ATM[26] and AutoML-gs have been evaluated. In order to analyse the competency of the AutoML framework itself, pure automated approaches were used with no manual interventions in feature engineering, data cleaning, data coding or parameter optimisation processes.

### 4.1 Description of Machine Learning methods

Different machine learning approaches have been grouped broadly.

#### 4.1.1 Gaussian Based Classification

Gaussian process classifier or GP Classifier[41] considers observations to be drawn from a Bernoulli distribution with probabilities relating to latent functions via sigmoid. Probit transformations define the likelihood model as a function of gaussian cumulative distribution function. In order to account for the non-linearities in the latent space, the covariance function is defined as linear sum of constant, linear, and Matern composite kernels.

#### 4.1.2 Feature Trimming Method

The feature trimming method or FT method[33] implements feature set reduction by introducing a random vector in the feature set, calculating the feature importance based on relative mean decrease of Gini-index(derived from random forests) thereby eliminating features scoring less than the dummy variable. Support Vector machines with Gaussian Kernel (RBF-SVM) are further training on the resulting features.

#### 4.1.3 Distance Weighted Discriminant method

The distance weighted discrimination(DWD) method[17] is a single base classifier where the high dimension low sample size(HDLSS) property is a subspace in the whole feature space. The HDLSS settings are known to work well with generative statistical models that better describe similarity between unobserved multimodal groups. Hence, the linear projection vectors with the data-piling property classify the data and the approach combined with Support Vector machines(SVM) entails accounting for distances from sample to separating hyperplane by associating a misclassification penalty cost C (=300) to sample points away from the hyperplane. The DWD method also further uses a second-order cone programming(SOCP) optimisation.

#### 4.1.4 Other methods

Popular methods include -

i. K nearest neighbours ^3^
ii. Multiple linear support vector instances^4^ amongst others.

### 4.2 Automated Machine Learning methods

The description and evaluation of specified tools show that while automating strenuous aspects of the pipeline can enhance developer productivity, some tools are not yet capable of catering to tasks with little data, or to those with limited domain knowledge.

#### 4.2.1 Ludwig

Ludwig is a Tensorflow based, deep-learning, encoder-decoder based toolbox that eliminates the need to write any code. Under the hood, Ludwig performs model training on Horovod[28], open source distributed training framework. Ludwig can generalise across training set types but for structured data, dependencies involve YAML files which allow control over model parameters. Intermediate pre-processed hdf5 file is built along with meta-data that maps input tensors to corresponding labels. During prediction, metadata is loaded to mimic the mapping of new data to its label thereby preventing retraining and re-computation of tensor values. This system of encoding-decoding works well here as the structure of the train data is very similar to that of the test data. The combiner between the encoder-decoder combines features into a single representation.

**Figure 5:**
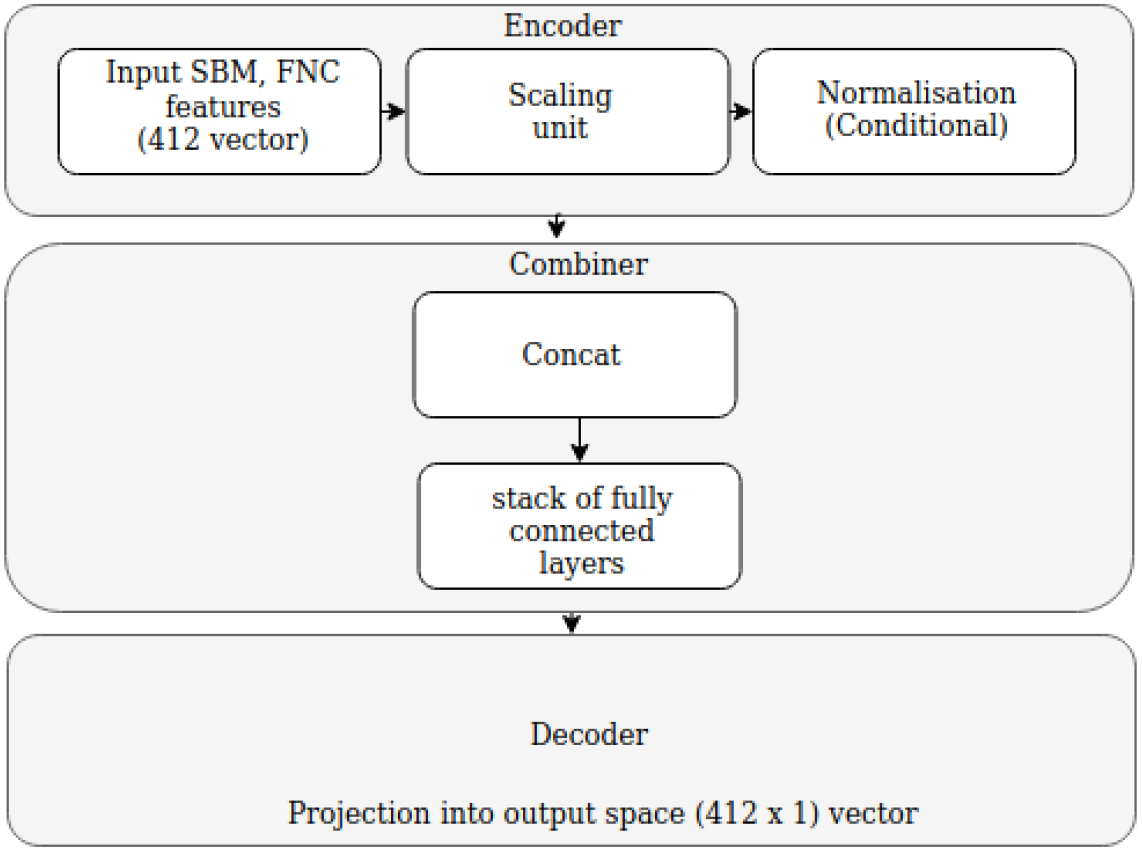
Architecture of Ludwig^5^

##### (a) Default Configuration

Features are automatically converted to a 412-vector and the default model with one encoder(stacked CNN model), one decoder(potentially empty stack of fully connected layers) and one combiner(concatenation by default) finally projects onto a single prediction. The non tensor inputs are reduced to their sums and the parameters by default are: mean squared error for loss and ReLU activation function with no dropout. Training runs for 100 epochs with Adam optimiser (beta1: 0.9, beta2: 0.999, epsilon: 1e-08), learning rate: 0.001 and decay rate of 0.96.

##### (b) Other Configurations

Ludwig does not directly support ensemble modeling. Experimentation with deep learning modules including:

i. Increasing the number of layers,
ii. Increasing layer size (from 256 to 512),
iii. Input normalisation of SBM and FNC features using z-score or minimax,
iv. Other activation functions like sigmoid and
v. Dropout regularisation

but these did not lead to substantial performance enhancement.

#### 4.2.2 H2O

An in-memory distributed scalable machine learning platform, H2O.ai finds an open port and connects localhost to a H2O cluster instance. The internal architecture of H2O is as illustrated in Figure 6.

**Figure 6:**
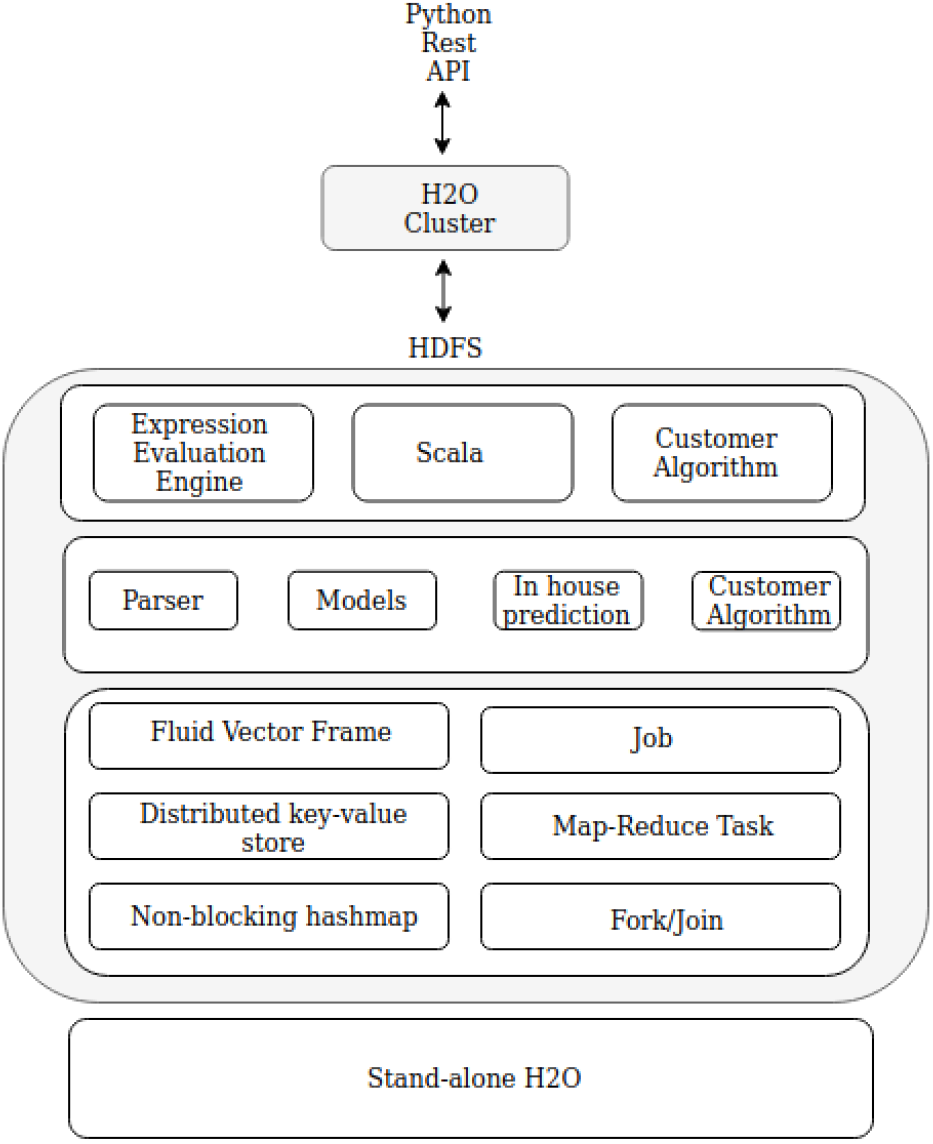
Architecture of H2O^6^

##### (a) Default H2O.ai Configuration

The default H2O configuration accounts for the input data via 5 fold cross validated model performance. The framework performs neural architecture search optimising for the AUROC metric and ranks all obtained algorithms on a leaderboard as per training performance which in this case includes: gradient based methods, stacked ensembles, generalized linear models, distributed random forests and XGBoost frameworks respectively.

##### (b) H2O.ai deeplearning

H2O’s deep learning module is based on a multi-layer feed-forward artificial neural network (multilayered perceptron) trained with stochastic gradient descent using back propagation. Each computational node trains a copy of the global model’s parameters with asynchronous multi-threading and contributes across the network through periodic model averaging. The different experimentation settings with the deep learning module include:

i. Activation functions - tanh, ReLU, maxout
ii. Regularisation functions - dropout
iii. Categorical encoding methods - one hot internal encoding: N+1 new columns for categorical features with N levels, and sort by response encoding: reordering of levels according to mean response.

##### (c) Other configurations

Ensemble and stacking did not yield substantial results.

#### 4.2.3 ATM - Autotune models

This multi-user, multi-data AutoML framework allows an end-to-end automated neural pipeline to search for the best model and henceforth train the best found algorithm like a normal sklearn model which is then pickled to classify on new data. The architecture of ATM is as follows:

**Figure 7:**
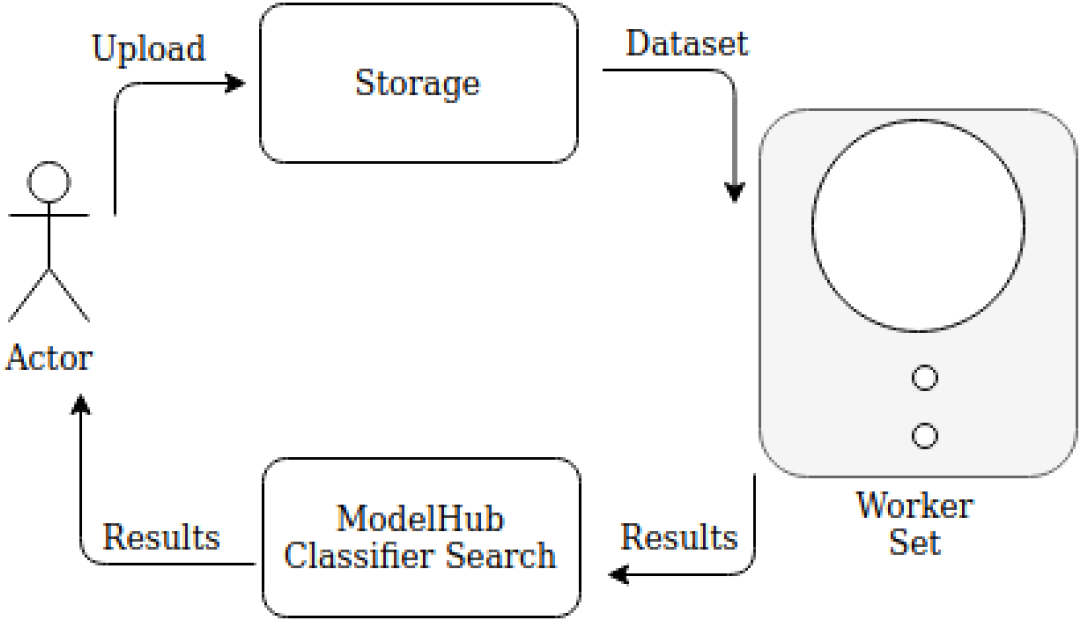
Architecture of ATM[26]

##### (a) Default Configuration

In the default configuration, search space of 100 classifiers is explored. The best performing model is a K-Nearest Neighbor(KNN) with the ball tree algorithm(k=42). Default models in queue include: logistic regression, decision trees and variants of KNN.

##### (b) Other Configurations

Increasing the budget of the classifiers leads to exploration of variants of KNN with different k values but does not necessarily lead to better performance on the private set. Using other methods like support vector machines (SVM), stochastic gradient descent (SGD), Gaussian and multinomial naive Bayes (NB), random forests, multi-layer perceptrons (MLP) and boosting does not improve performance significantly.

#### 4.2.4 AutoGluon-Tabular

This automated framework offers multi-layer ensembling that is trained layer-wise to translate raw data into predictions. Skip connections in stack ensembling and neural network embedding as well as repeated k-fold bagging are employed to curb overfitting. AutoGluon-Tabular is also capable of inferring properties of the prediction task like variable types and problem class. The architecture of AutoGluon-Tabular is as illustrated in Figure 8.

**Figure 8:**
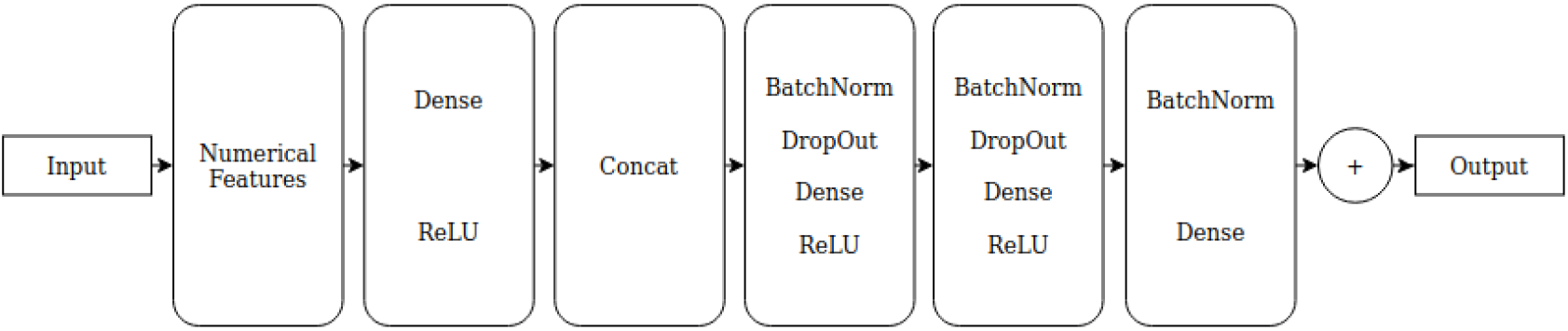
Architecture of AutoGluon[24]

##### (a) Default Configuration

AutoGluon by default enables prototyping of deep learning systems with automatic hyperparameter tuning, model selection and data processing while choosing a random training-validation data split with conditional stratified sampling strategies. Since the described methods are computationally intensive, the process is parallelised across threads and machines (depending on amount of available distributed resources). However, AutoGluon opts for tuning on validation data using internal knobs and hence, internal performance estimated maybe higher as compared to private or public AUC. Predictions are auto-optimised to reduce inference latency.

##### (b) Other configurations

Bagging, stacking or ensembling did not lead to substantial performance enhancements.

#### 4.2.5 ML Box

MLBox is an automated machine learning framework that uses drift scores to select important feature attributes by discarding variables automatically that have distributions that do not align with the rest of the data(covariate shift) and therefore, do not contribute to the prediction process which in turn also enhances performance on biased data by relating the ROC score as measure of drift. If the drift is high, then the sets of data are discernible and if low, they are not. MLBox implicitly uses a drift thresholder that retains only features lesser than the threshold. The architecture is as shown in Figure 9.

**Figure 9:**
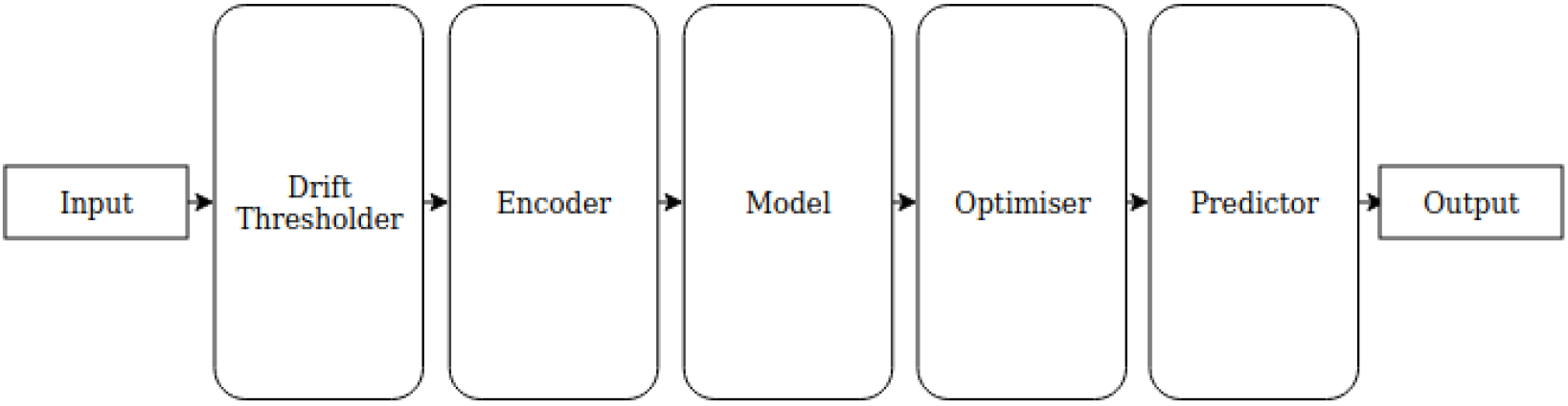
Architecture of ML Box^7^

##### (a) Default configuration

There is no default configuration of MLBox for the space search. So, the following space configurations were evaluated:

i. A stacking classifier with multiple Adaboost classifiers
ii. Encoding strategy with random projections, label encodings and entity embeddings. Feature selection threshold with uniform search in the [0.001, 0.2] space. Final estimation strategy in the random forests, extra trees and light gradient based methods space.

##### (b) Other configurations

Widening the search space does not necessarily lead to better private AUC performance.

#### 4.2.6 AutoML-gs

AutoML-gs^8^ generates raw Python code using Jinja templates and trains the statistical model in a subprocess by trying differentiating hyperparameters to find the best model. It automatically infers feature types and then uses a ETL strategy for feature optimisation according to their hyperparameters. Encoders used are stored as serialised JSON files with consistent, discrete hyperparameter values instead of uniform hyperparameter values to accurately gauge tuning impact. The architecture is as described in Figure 10.

**Figure 10:**
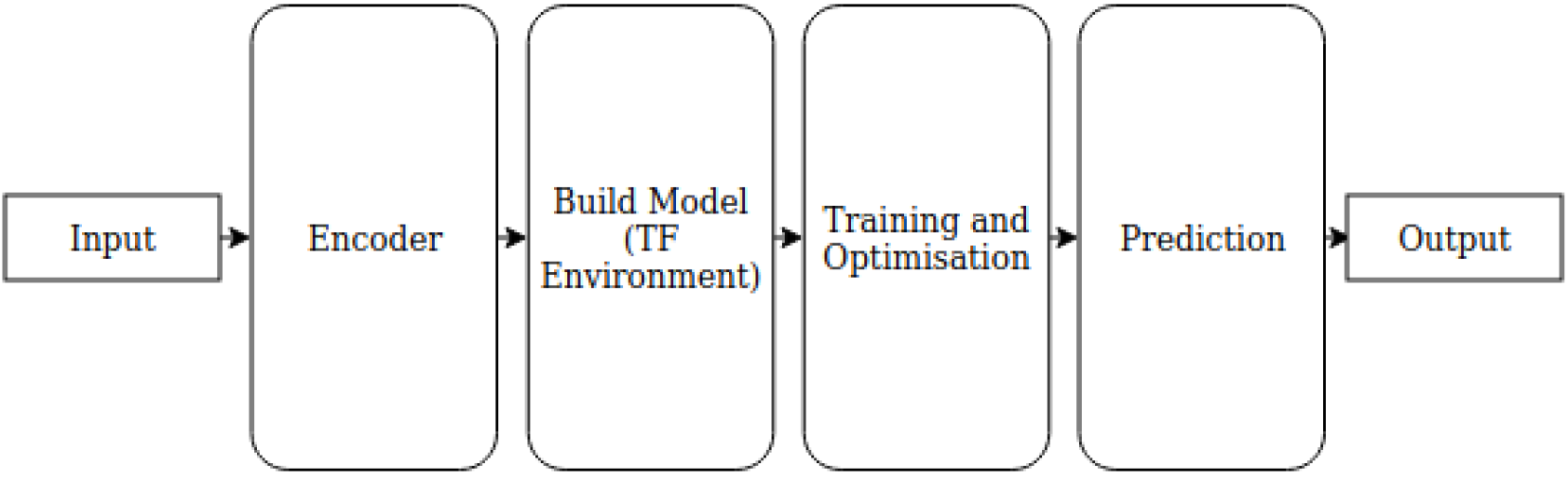
Architecture of AutoML-gs^9^

##### (a) Default Configuration

There are two environments available in AutoML-gs: Tensorflow and XGBoost. Experimentation has been carried out on both here.

## 5 Results

The results from the mentioned models and configurations are evaluated broadly based on the “area under the receiver operating characteristics curve” (AUROC) statistic under the following subcategories. The ROC represents a probabilistic curve, and AUC represents degree of separability. Higher the statistic, better the model is at segregating schizophrenics against non-schizophrenics.

The tools and frameworks are compared on the following parameters for the best performing configuration within each tool: (a) Public AUC which involves evaluation of about one percent of the given data. (b) Private AUC which involves evaluation of about 99 percent of (hidden) test data. (c) Overall AUC which involves evaluation on all test data. (d) Model Stability and Robustness (Model-SR) which is defined based on private and public AUC. (e) Interpretability.

It was observed that the overall AUC led to more reasonable assessments about model quality than any other metric[27]. Note: Since private data involves most of test data, public data has not been a priority for performance or optimisation.

The default configuration mode in each tool performs as tabulated in Table 1.

**Table 1:**
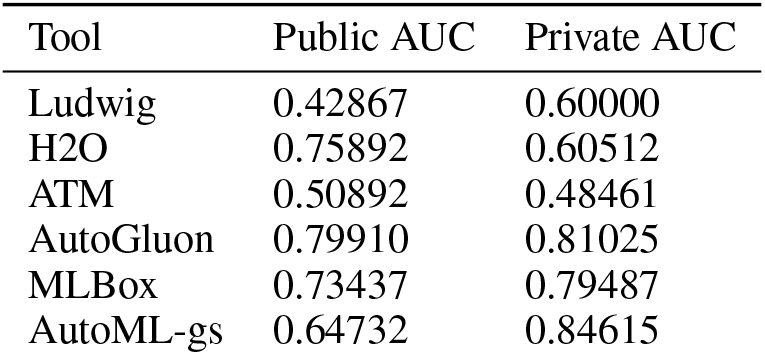
Comparison of Default configurations of AutoML methods

The visual analysis of the data is as shown in Figure 11.

**Figure 11:**
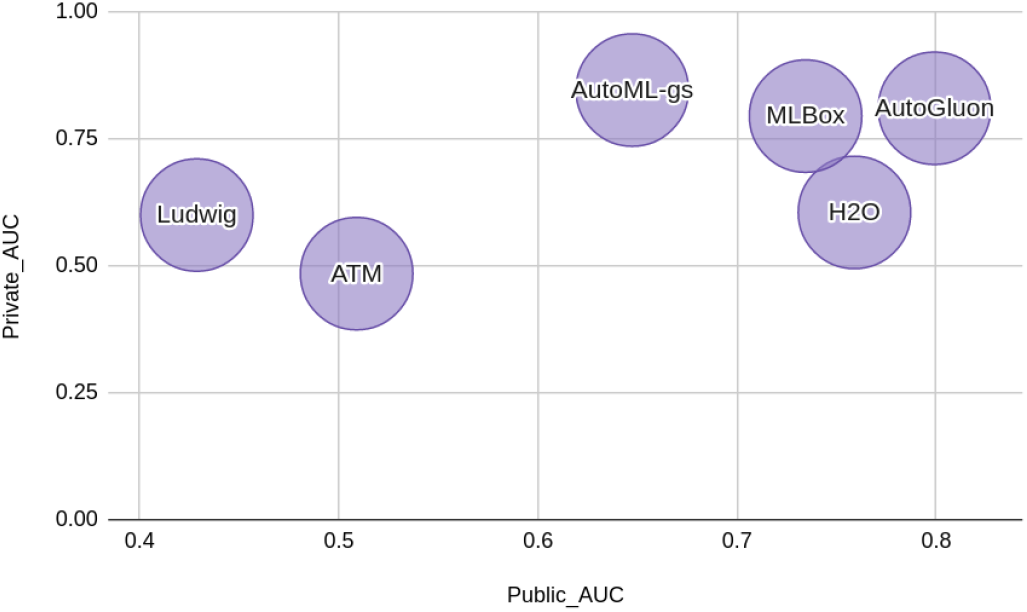
Relative performance of AutoML frameworks: Default configuration

The performance of 2087 entries from 291 participants[27] has been aggregated into the visualisation below in terms of public and private AUC.

**Figure 12:**
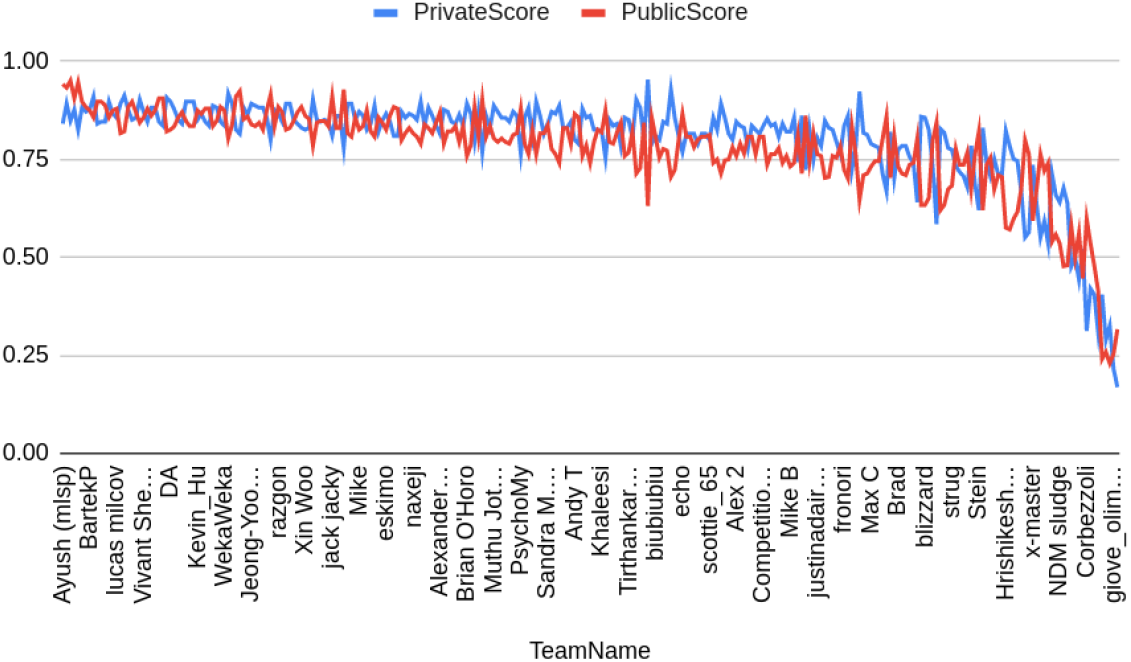
Relative performance of ML frameworks

### 5.1 Public AUC

Most of machine learning methods perform well on private data, but do not perform nearly as well on the public data with the exception of SVM.

**Table 2:** Relative Performance of Machine Learning methods: Public AUC

| Method | Public AUC |
| --- | --- |
| GP Classifier | 0.70536 |
| FT Method | 0.64732 |
| DWD Method | 0.84375 |

However, the best performance on the public data is 0.95758 and the highest scoring AutoML model - one hot encoding of categorical features with rectifier on H2O - is 0.95384. This score has not been optimised further here as our obtained performance is already at the top 1 percent of all models.

**Table 3:** Relative Performance of AutoML methods: Public AUC

| Method | Sub-configuration | Public AUC |
| --- | --- | --- |
| Ludwig | 512 FC layers | 0.55357 |
| H2O | ReLU activation | 0.88839 |
| H2O | sortByResponse encoding with dropout | 0.80803 |
| H2O | OneHotInternal encoding with rectifier | 0.95384 |
| AutoGluon | Default | 0.79910 |
| MLBox | Default | 0.73437 |

The performance of all models is as shown in Figure 13 where the y-axis represents the Public AUC score and x-axis represents the model. H2O SR refers to the H2O sort by response encoding with dropout configuration and H2O 1HiE refers to the H2O one hot internal encoding with rectifier configuration.

**Figure 13:**
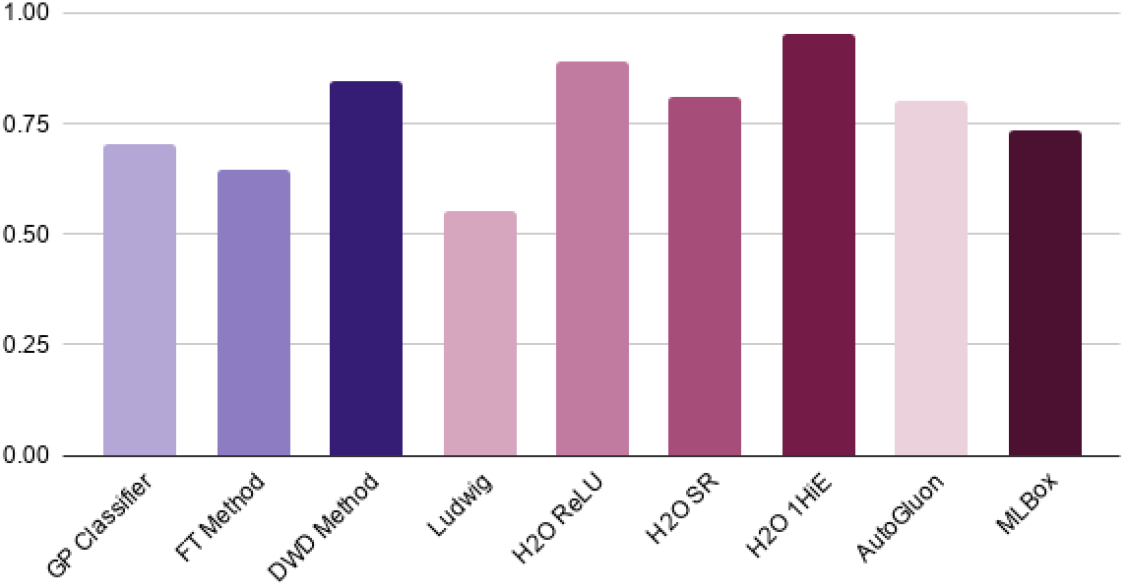
Relative Performance of ML and AutoML methods: Public AUC

#### 5.1.1 Private AUC

Gradient based methods seem to perform the best in terms of generalisation as they account for intricacies including short-scale, non-linearities in the latent space while maintaining model flexibility. It is notable that - most of these automated machine learning tools lean towards gradient based classifiers similar to developer behavior[27].

**Table 4:**
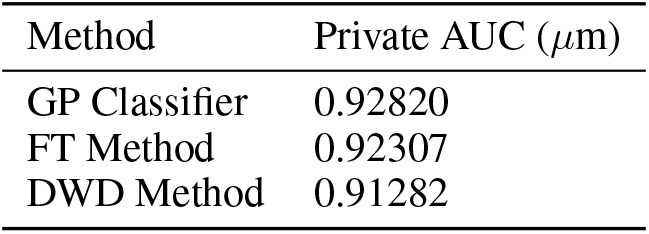
Relative Performance of Machine Learning methods: Private AUC

| Method | Private AUC ($\mu m$) |
| --- | --- |
| GP Classifier | 0.92820 |
| FT Method | 0.92307 |
| DWD Method | 0.91282 |

**Table 5:** Relative Performance of AutoML methods: Private AUC

| Method | Sub-configuration | Private AUC |
| --- | --- | --- |
| Ludwig | Default | 0.60000 |
| H2O | Gradient-based | 0.72307 |
| ATM | Classifiers=500 | 0.83333 |
| AutoGluon | Default | 0.6923 |
| MLBox | Default | 0.73437 |
| AutoML-gs | Default | 0.82820 |

**Figure 14:**
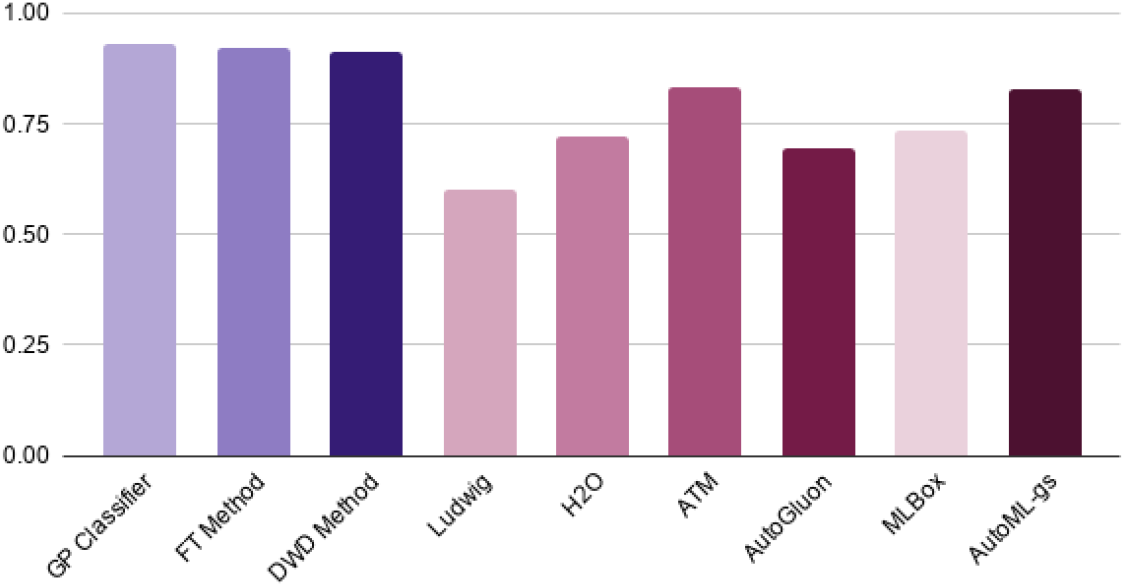
Relative Performance of ML and AutoML methods: Private AUC

On further experimentation with H2O’s deep learning module configurations, the following observations were made:

a. ReLU generalised better with dropout regularisation.
b. Models with categorical encoding tended to perform best with respect to Private AUC.

**Table 6:**
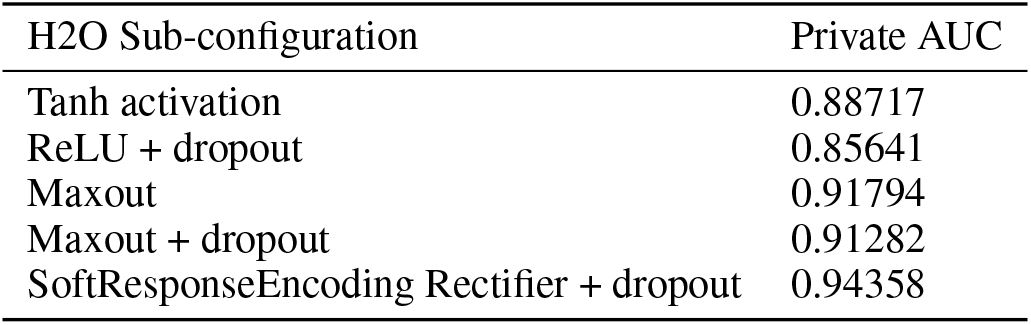
Relative Performance of H2O deep learning configurations: Private AUC

| H2O Sub-configuration | Private AUC |
| --- | --- |
| Tanh activation | 0.88717 |
| ReLU + dropout | 0.85641 |
| Maxout | 0.91794 |
| Maxout + dropout | 0.91282 |
| SoftResponseEncoding Rectifier + dropout | 0.94358 |

H2O deep learning configured models perform comparably to ML models.

**Figure 15:**
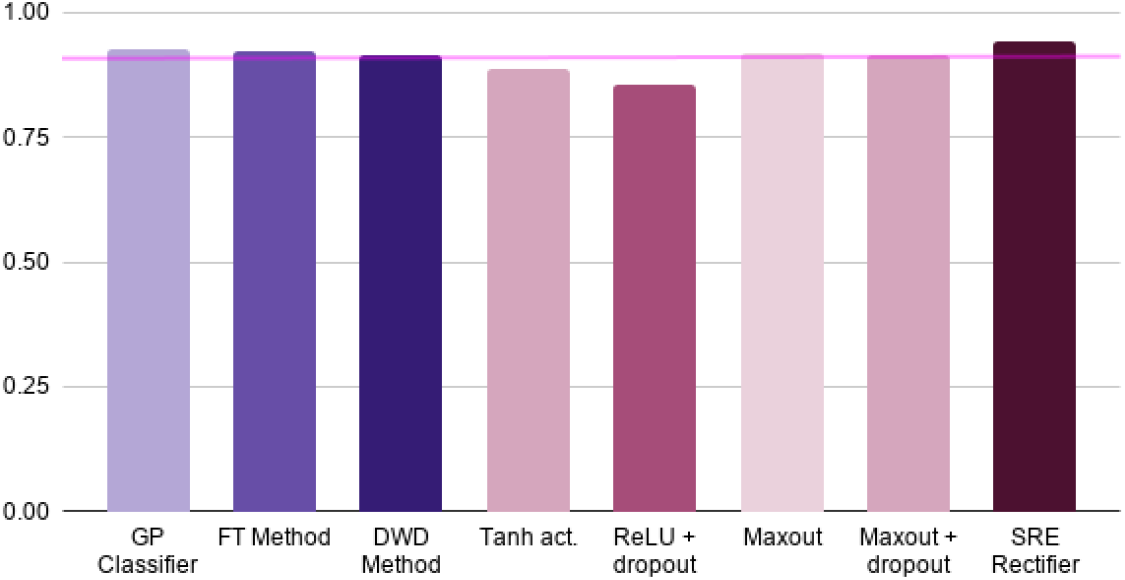
Relative Performance of H2O and ML methods: Private AUC

#### 5.1.2 Overall AUC

The Overall AUC helps favor consistent models while ensuring retention of generalisability. However, none of the machine learning models have an Overall AUC of >0.9[27] which is hereby known as “the 0.9 wall”. The best models in this respect average to about 0.88 but this is an important reinforcement of the performance of machine learning classifiers in challenging settings. It is also worthwhile to consider the fact that although this result is obtained with features derived from independent components using spatio-temporal regression highlighting the predictive utility of imaging data, the identification of appropriate predictive features within the entire space of features is still a challenging process.

**Figure 16:**
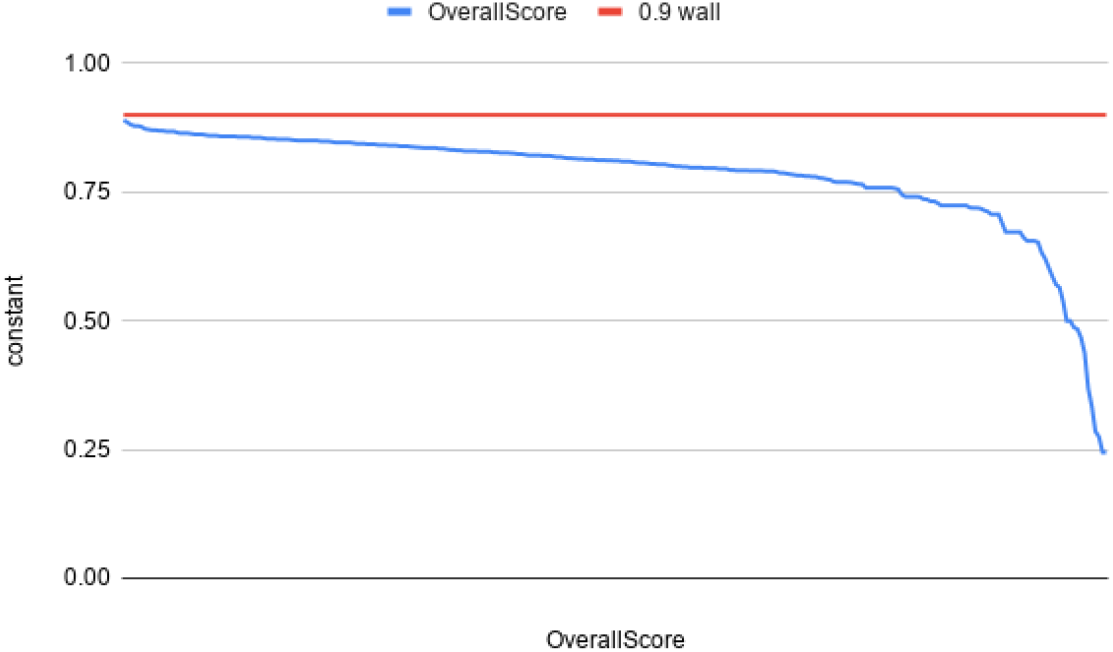
Relative performance of all machine learning models(290): Overall AUC

Most machine learning methods work better than the benchmark average with SVM based algorithms performing the best.

**Table 7:**
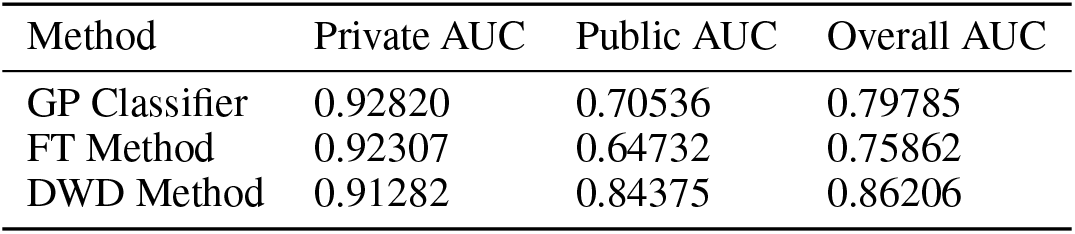
Relative Performance of Machine Learning methods: Overall AUC

| Method | Private AUC | Public AUC | Overall AUC |
| --- | --- | --- | --- |
| GP Classifier | 0.92820 | 0.70536 | 0.79785 |
| FT Method | 0.92307 | 0.64732 | 0.75862 |
| DWD Method | 0.91282 | 0.84375 | 0.86206 |

AutoML tools automate the process of feature identification efficiently even with little training data and high dimensional features. Table 15 highlights that subtle configuration changes such as changing categorical encoding methodology or activation functions can comfortably scale the 0.9 wall.

**Table 8:** Relative Performance of AutoML methods: Overall AUC

| Method | Private AUC | Public AUC | Overall AUC |
| --- | --- | --- | --- |
| H2O with activation | 0.91794 | 0.88839 | 0.9 |
| H2O encoding with ReLU activation | 0.94358 | 0.95384 | **>0.9** |

The aggregate Overall AUC of the data is as shown.

**Figure 17:**
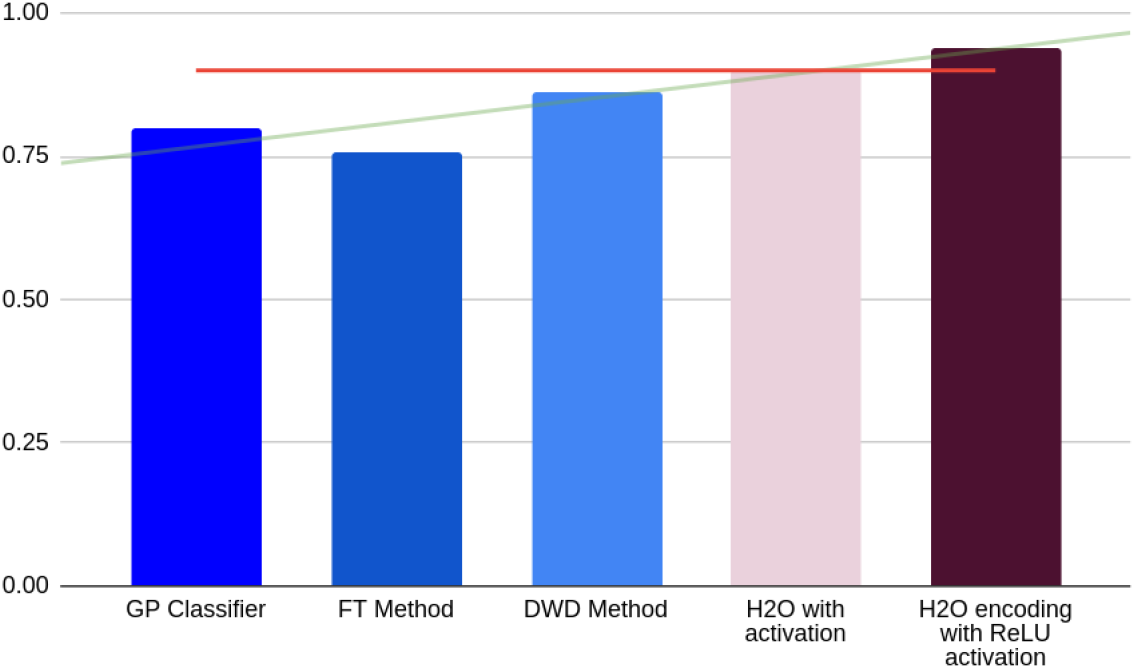
Relative Performance of ML and AutoML methods: Overall AUC

#### 5.1.3 Model Stability and Robustness(Model-SR)

Model stability and robustness which we define here as Model-SR is:

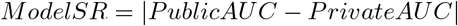

and a model is said to be stable if this absolute difference is nearly 0. However this does not seem to be the case with many machine learning models. The Model-SR calculated from 290 models is as shown in Figure 18.

**Figure 18:**
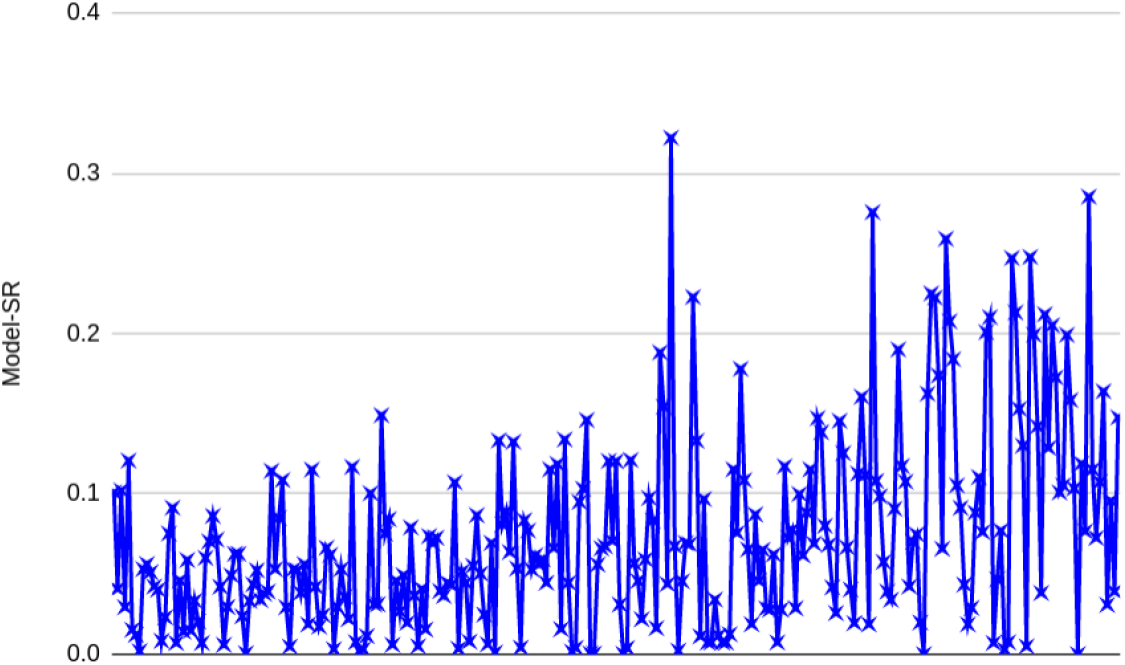
Relative performance of ML models (290): Model-SR

Amongst the models that perform best with respect to private AUC, Gradient based methods show the highest degree of variability with Model-SR as even observed with Gradient based models from Automated tools. Support vectors models display maximum stability.

**Table 9:**
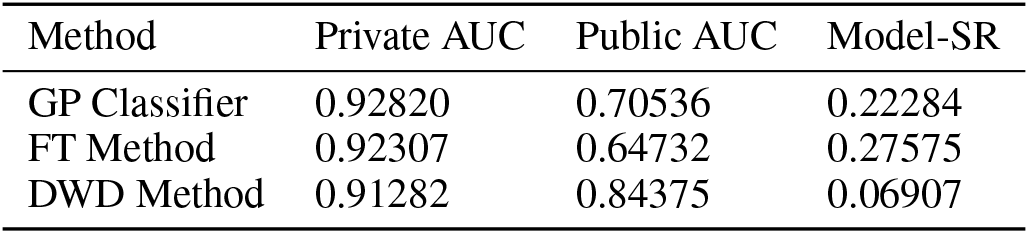
Relative Performance of Machine Learning methods: Model-SR

| Method | Private AUC | Public AUC | Model-SR |
| --- | --- | --- | --- |
| GP Classifier | 0.92820 | 0.70536 | 0.22284 |
| FT Method | 0.92307 | 0.64732 | 0.27575 |
| DWD Method | 0.91282 | 0.84375 | 0.06907 |

The effect of stability of automated tools is observed from 132 different configurations of more than six different tools on multimodal schizophrenia data. The most stable AutoML models are AutoGluon followed by MLBox and H2O (default configuration). Deep learning H2O modules with eigen encoding and maxout display the maximum stability at 0.01921 while AutoML-gs and ludwig seem to display the least stability.

**Table 10:** Relative Performance of AutoML methods: ModelSR

| Tool:Configuration | Difference AUC |
| --- | --- |
| AutoGluon: default | 0.00927 |
| MLBox: default | 0.0652 |
| ATM: 500 classifier<sub>budget</sub> | 0.10119 |
| H2O: default | 0.0445 |
| H2O: activation | 0.18 |
| H2O: ReLU + dp | 0.18384 |

**Figure 19:**
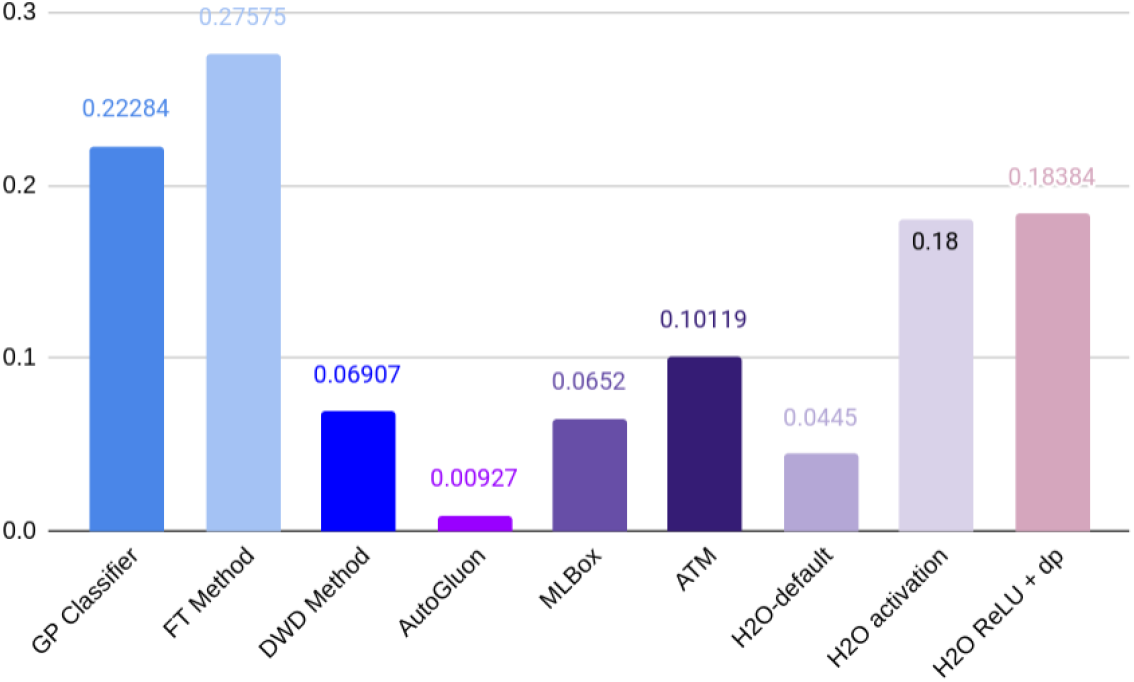
Relative performance of ML vs AutoML models: Model-SR

#### 5.1.4 Interpretability

AutoML tends to further obscure black-boxed models. But interpretability is of utmost value for healthcare diagnostics and for other safety critical applications in general. While most of the tools used in the study enable external augmentations and interpretable enhancements, few internally support interpretability.

##### MLBox

MLBox has an internal prediction feature that highlights the top features as per their drift coefficients that most influence the output.

**Figure 20:**
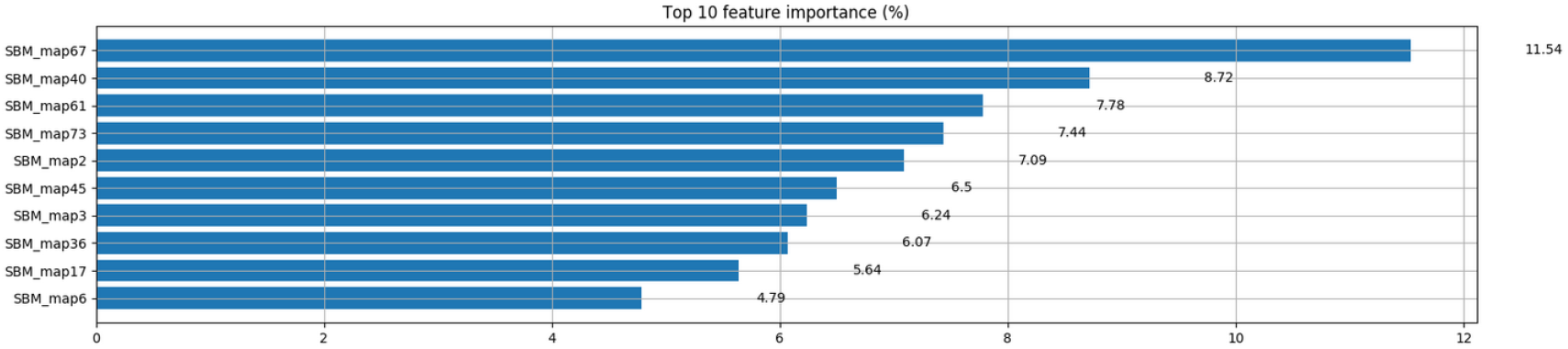
Relative Feature importance in MLBox

##### AutoML-gs

The output predictions are shipped along with json files containing the mean, scale and variance of every feature in the dataset that the encoder model is trained on. It also contains additional information about the training time taken, the performance parameters including - log loss, accuracy, AUC, precision, recall and f1 score for every epoch. The model weights along with the entire pipeline can be accessed and manipulated. This makes this AutoML framework nearly equivalent to any machine learning framework.

##### H2O

In case of deep learning, hidden model parameters, variable importance, graphs, weights/biases, and the status of neuron layers namely - layer number, units, type, dropout, L1 regularisation, L2 regularisation, mean rate, rate root-mean-square, momentum, mean weight, weight root-mean-square, mean bias, and bias root-mean-square are logged. Visualisations can be built externally with Tableau.

##### Ludwig

Since the definitions are externally specified through the YAML structure, interpretability with respect to feature engineering lies much outside the framework than within. Experimental runs are saved along with predictions, probabilities, and different classes of statistics including: (A) Class statistics which includes confusion matrix, average precision across samples and (B) Performance related metrics including loss, accuracy, F1, kappa score, token accuracy, precision, and recall (C) Per class statistics for all True/False values - fallout, discovery rates, false negative rates, omission rates, false positives, hit rate, informedness, markedness, matthew’s correlation coefficient, miss-rate, predictive values, specificity, sensitivity, true negative rate, and true positive rate. It also accounts for logging model hyperparameters for each feature, indexing, checkpointing, tracking training progress, and working with meta data. These parameters can be visualized externally via Tensorboard.

##### ATM

Each of the models and model parameter metrics in the range of the classifier budget are saved for later access. The framework is nearly equivalent to other machine learning models in terms of interpretability.

##### AutoGluon

Models generated are saved along with information about existing and generated features. On running this instance, the dask worker space is first locked before start of training. Visualizations can built externally via tensorboard or mxboard.

## 6 Conclusion and Scope for future work

Automated machine learning tools have a remarkable ability to effectively deal with multimodal data under high dimension low sample size settings(HDLSS) as demonstrated. The space of configurations and frameworks has transformed over the years but the generalisability of these methods to raw or lightly processed data and their ability to automatically identify features from whole brain information in a clinical setting still remains a challenge. With respect to healthcare systems in particular, there is a need to develop automated frameworks that are not only consistent across executions for reproducibility, but also more transparent, ethical, and interpretable. Future research in this direction would hence require tools that better understand data, and hence are able to employ reasonable choice of algorithmic models along with correspondingly different tuning strategies.

## 7 Acknowledgements

i. Data collection (partially described in [19]), available on Kaggle^10^, was performed at the Mind Research Network, and funded by a Center of Biomedical Research Excellence (COBRE) grant 5P20RR021938/P20GM103472 from the NIH to Dr. Vince Calhoun.
ii. The suggestions of Erin LeDell (H2O.ai), @PGijsbers(gama), Piero Molino (Ludwig), Dr. V Krishnamurthy, Dr. S Natarajan, Anil B Murthy and Alex De Romblay (MLBox).

## Footnotes

0 Accepted as workshop contribution at Women in Machine Learning (WiML) co-located with ICML 2020. Related code can be found at: https://github.com/GaganaB/AutoML

1 https://www.health.harvard.edu/a_to_z/schizophrenia-a-to-z

2 www.fil.ion.ucl.ac.uk/spm/software/spm5

3 https://github.com/KonstantinTogoi/MLSP2014

4 https://github.com/gabegaster/kaggle_schizophrenia_2014

5 https://uber.github.io/ludwig/

6 http://docs.h2o.ai/h2o/latest-stable/h2o-docs/architecture.html

7 https://mlbox.readthedocs.io/en/latest/

8 urlhttps://mljar.com/blog/automl-software-list/

9 https://github.com/minimaxir/automl-gs

10 https://www.kaggle.com/c/mlsp-2014-mri/

